# Self-Supervised Learning Improves Accuracy and Data Efficiency for IMU-Based Ground Reaction Force Estimation

**DOI:** 10.1101/2023.10.25.564057

**Authors:** Tian Tan, Peter B. Shull, Jenifer L. Hicks, Scott D. Uhlrich, Akshay S. Chaudhari

## Abstract

**Objective:** Recent deep learning techniques hold promise to enable IMU-driven kinetic assessment; however, they require large extents of ground reaction force (GRF) data to serve as labels for supervised model training. We thus propose using existing self-supervised learning (SSL) techniques to leverage large IMU datasets to pre-train deep learning models, which can improve the accuracy and data efficiency of IMU-based GRF estimation.

**Methods:** We performed SSL by masking a random portion of the input IMU data and training a transformer model to reconstruct the masked portion. We systematically compared a series of masking ratios across three pre-training datasets that included real IMU data, synthetic IMU data, or a combination of the two. Finally, we built models that used pre-training and labeled data to estimate GRF during three prediction tasks: overground walking, treadmill walking, and drop landing.

**Results:** When using the same amount of labeled data, SSL pre-training significantly improved the accuracy of 3-axis GRF estimation during walking compared to baseline models trained by conventional supervised learning. Fine-tuning SSL model with 1–10% of walking data yielded comparable accuracy to training baseline model with 100% of walking data. The optimal masking ratio for SSL is 6.25–12.5%.

**Conclusion:** SSL leveraged large real and synthetic IMU datasets to increase the accuracy and data efficiency of deep-learning-based GRF estimation, reducing the need for labeled data.

**Significance:** This work, with its open-source code and models, may unlock broader use cases of IMU-driven kinetic assessment by mitigating the scarcity of GRF measurements in practical applications.

## I. Introduction

**B**IOMECHANICAL assessment of human movement is important for ascertaining human health. Kinetics during locomotion can be used to predict injury risk and assess rehabilitation outcomes. For example, ground reaction forces (GRF) during a sit-to-stand task can provide insights into rehabilitation outcomes after hip injury [1], and knee adduction and flexion moments during walking have been associated with knee osteoarthritis [2]–[4]. Additionally, GRF patterns during drop landing, where participants step off an elevated platform and land on the ground, are associated with several lower-extremity injuries [5], [6].

Despite the value of estimating kinetics, measurement of GRF and joint moments typically requires gold standard measurements from force plates and marker-based motion capture. However, these devices are expensive, require trained personnel to operate, and confine the measurement environment. Consequently, such challenges associated with acquiring kinetic data limit the accessibility of biomechanical assessment and prevent translation of biomechanical studies to larger and diverse populations.

Using wearable sensors to monitor kinetic parameters in natural environments represents a promising opportunity for moving disease characterization, prevention, and rehabilitation assessment into real-world settings [7]. Inertial measurement units (IMUs) are a practical choice for out-of-lab assessment due to their low cost and small form factor. IMUs typically measure acceleration and angular velocity (and magnetic field in some scenarios), which can directly be used to derive body segment orientation for subsequent kinematic estimation.

Prior studies have combined physics-based models and optimization methods to estimate kinetics from IMU data [8]–[10] However, optimization methods require a relatively high computation time and depend on task-specific movement objectives (e.g., minimizing energy, maximizing speed) [11], [12]. Apart from physics-based models, other studies have attempted to employ deep learning models that directly map IMU data to kinetic parameters in an end-to-end manner [13]– [15]. To train these models, previous studies have collected synchronized IMU, GRF, and marker-based motion capture data during hundreds of minutes of walking and running on treadmills equipped with force plates. The GRF and marker data then serve as “labels” to train models using conventional supervised learning. However, collecting these “labels” in large quantities to train accurate deep learning models is not always feasible for motions beyond treadmill gait. For overground walking, since GRF data is typically collected via one to three floor-embedded force plates, each trial often yields only one or several steps with “labels”. The scarcity of data also holds true for a variety of other motions including sprinting, cutting, and jumping [16].

Self-supervised learning (SSL) may be used to pre-train deep learning models to mitigate the challenge of data scarcity [17]–[19] It may also improve models’ robustness to noisy data [20] and distribution shifts where the distribution of model training data differs from that of testing data [21]. SSL aims to pre-train deep learning models by designing model pre-training tasks that are solely based on “unlabeled” data. One widely used pre-training task is masking and reconstruction, where a random portion of the input data is masked and the model is trained to learn the data representation by reconstructing the masked portion. When limited “labeled” data is available for a downstream prediction task of interest, representations learned from pre-trained models can enhance the training process, and lead to better performance than conventional supervised learning with randomly initialized parameters. SSL with masking and reconstruction has facilitated the training of powerful language models [17], [22], vision models [18], and multi-modal models [19]. However, it is unknown whether SSL with masking and reconstruction can learn IMU representations that improve performance on downstream tasks of estimating kinetics. Additionally, unlike the fields of natural language and vision, the optimal pretraining dataset and strategy for this IMU-based application remain unclear.

The aim of this paper is to use existing SSL techniques to improve the accuracy and data efficiency of an IMU-driven deep learning model, and open source it on GitHub. We first investigated the optimal SSL strategy using a large “unlabeled” dataset of 500 participants with real and synthetic IMU data. We then compared the performance and data efficiency of SSL pre-trained models on estimating GRF compared to baseline models whose parameters are randomly initialized following conventional supervised learning. We first estimated GRF on overground and treadmill walking datasets, which was within the distribution of the SSL pre-training datasets. Then, we estimated GRF on a drop landing dataset, which is not present in the pre-training dataset. To facilitate broader usage and reproduction of work, we make our framework broadly usable by releasing our source code and trained models.

## II. Methods

### A. Pre-training Datasets

We used three datasets for our SSL pre-training: real IMU data [23], synthetic IMU data [24], and a fused dataset combining them both (Table I). The real IMU dataset has 17 IMUs, and we used eight of them that were placed on the trunk and pelvis, as well as both thighs, shanks, and feet. For the synthetic IMU dataset, we generated data on the same eight body segments. Each IMU has a 3-axis accelerometer and a 3-axis gyroscope. The z-axes of the IMUs were aligned with the segment surface normal, x-axes were pointing left during standing, and y-axes were perpendicular to the x- and z-axes following the right-hand rule (Fig. 1).

**TABLE I.** Two datasets for SSL pre-training and three datasets for downstream evaluation.

| Type | Ref. (name) | Motion | # of Participants<sup>a</sup> | # of Windows <sup>a, b</sup> |
| --- | --- | --- | --- | --- |
| Pre-training Datasets | [23] (MoVi) | 21 motions including walking | 90 | 14,418 |
|  | [24] (AMASS) | 100+ motions including walking | 500 | 713,828 |
| Downstream Datasets | [25] | Overground walking | 19 | 510 |
|  | [26] | Treadmill walking | 17 | 23,481 |
|  | [27] | Drop landing | 15 | 509 |

<sup>a</sup> The pre-training datasets have more participants as well as comparable or more windows than downstream evaluation datasets.
<sup>b</sup> Each window is a model input segmented from original trials.

**Fig. 1.**
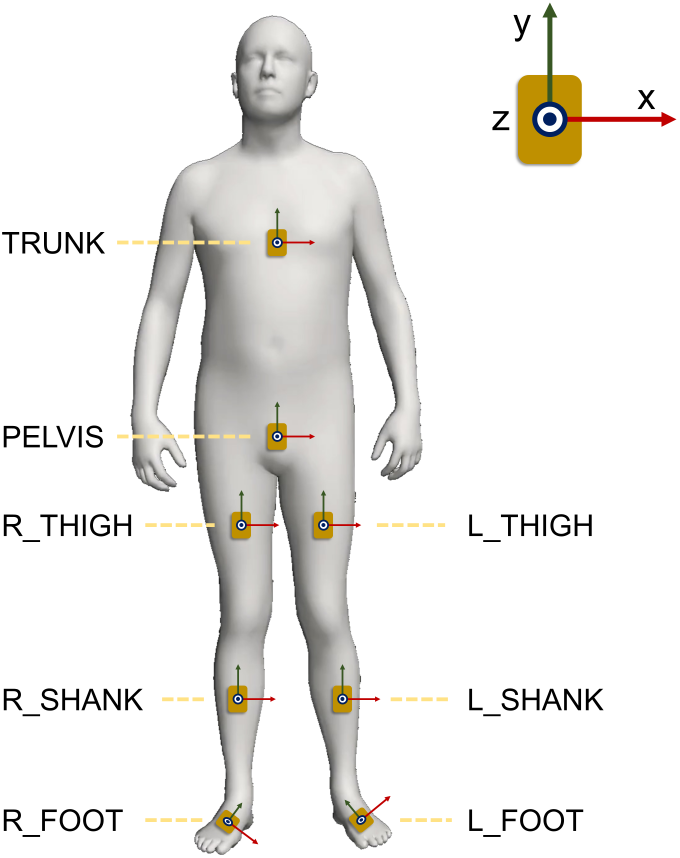
Locations and orientations of eight synthetic IMUs generated from the AMASS dataset. Red arrows, green arrows, and blue dots represent x-, y-, and z-axes of IMUs, respectively.

#### 1) Real IMU

This dataset is based on MoVi, a public human motion dataset with 90 participants who performed a collection of 21 everyday actions and sports movements such as walking, vertical jumping, and throwing [23]. Participants wore a 17-IMU suit that operated at 120Hz (Noitom, China). We linearly resampled each trial from 120Hz to 100Hz and then segmented it into windows of 128 time steps (1.28*s* each) with 50% overlap (0.64*s*) between successive windows. We excluded 18,760 (56.5%) windows in which participants were standing or sitting still, when the average standard deviation of all accelerometer axes was smaller than 0.2*m/s*^2^. In total, we obtained 14,418 windows and randomized their order for pre-training.

#### 2) Synthetic IMU

This dataset is based on AMASS, a large-scale, public human motion archive involving 500 participants [24]. AMASS unifies 23 existing motion capture datasets that have different marker sets, sampling frequencies ranging from 59–120Hz, and a variety of motions, with walking being one of the most common motions. Using kinematics to generate synthetic IMU data is a widely-used method for augmenting the volume of training data [28]–[31]. To generate synthetic IMU data, we low-pass filtered the joint angles and positions of each trial at 15Hz using a zero-lag, fourth-order Butterworth filter and resampled the filtered data to 100Hz. Subsequently, for each trial, we placed eight synthetic IMUs on their corresponding body segments (Fig. 1) five times with five different randomly generated orientation variations. Specifically, for each orientation variation, each of the eight IMUs is rotated by three random angles (within*±*10*deg*) along the x, y, and z axes, respectively. Then, we generated synthetic IMUs’ positions and orientations via forward kinematics using a prior pipeline [32]. Synthetic angular velocities were computed by taking the first derivative of orientation, whereas synthetic accelerations were computed by taking the second derivative of positions and incorporating gravity. Similar to the MoVi dataset pre-processing, we segmented the synthetic IMU data into windows of 128 time steps with 50% window overlap. We excluded 853,139 (54.0%) windows whose average acceleration was smaller than 0.2*m/s*^2^. We also excluded 12,408 (0.8%) windows that contained abnormally large synthetic acceleration or angular velocity with thresholds of 160*m/s*^2^ and 2000*deg/s*, respectively, which were determined based on measurement ranges of typical IMUs (MPU-9250, Invensense, USA and BHI-160, Bosch Sensortec, Germany). In total, we obtained 713,828 windows and randomized their order for SSL.

#### 3) Real and Synthetic IMU

This dataset combines real IMU data and synthetic IMU data described above, resulting in a total of 728,246 windows. Such a data dataset has the potential to leverage the benefits of a substantial data scale as well as authentic sensor noise representations.

### B. Downstream Datasets

We evaluated the performance of IMU representations from SSL pre-training in estimating GRFs for three different tasks, with corresponding datasets of synchronized IMU and GRF data: overground walking [25], treadmill walking [26], and drop landing [27]. The medial-lateral GRF (mlGRF), anterior-posterior GRF (apGRF), and vertical GRF (vGRF) of the right leg were used as the estimation target for all three tasks.

The IMU-to-segment position placements of downstream datasets do not need to be identical to those of the pretraining dataset (Fig. 1), as a prior study found that position variations within 100 mm may not substantially influence the accuracy of machine-learning-based GRF estimation models [29]. Meanwhile, for orientation placement, the real IMU dataset for pre-training has natural human-introduced placement variation while the synthetic IMU dataset for pretraining has*±*10*deg* orientation variations. Still, we assumed the following orientation placement convention: one of their surfaces aligned to the body surface and one of their edges aligned with the segment’s long axis. We manually verified whether the orientation placement conventions of each dataset are consistent with the assumed alignment. If not, we rotated IMUs along axes by multiples of 90 deg to match the assumed alignment.

#### 1) *Overground Walking*[25]

The original dataset contains 19 participants walking on stairs, ramps, treadmills, and level ground at various speeds, but we only used level-ground walking data to build the model to estimate GRF. The input IMU data are collected at 200Hz from four IMUs (Yost, USA) placed on the torso and right thigh, shank, and foot. GRF was measured by force plates (Bertec, USA) at 1,000Hz and low-pass filtered at 15Hz. We downsampled the synchronized IMU and GRF data to 100Hz to match that of the pre-training datasets. Each trial has three gait cycles with gold-standard GRF measurements. We extracted these gait cycles using windows of 128 time steps, starting at 40 samples before the stance phases. In total, there are 510 windows available for model training and evaluation.

#### 2) *Treadmill Walking* [26]

This dataset includes 17 participants walking on an instrumented treadmill with various speeds, foot progression angles, trunk sway angles, and step widths. Eight 8 IMUs (SageMotion, USA) were placed on the trunk, pelvis, and both thighs, shanks, and feet to collect data at 100Hz. The GRF of the right leg is measured by an instrumented treadmill (Bertec, USA) at 1,000Hz, low-pass filtered at 15Hz, and downsampled to 100Hz. For each trial, continuous walking data were segmented into windows of 128 time steps with 50% overlap. We retained all data windows because GRF is measured by treadmill-embedded force plates throughout the entire trial. In total, there were 23,481 windows available for model training and evaluation.

#### 3) *Drop Landing* [27]

This dataset includes 15 participants performing double-leg drop landing trials from a 30-cm-high box with various toe-out angles. Eight IMUs (Xsens, The Netherlands) were placed on the trunk, pelvis, and both thighs, shanks, and feet to collect data at 100Hz. The GRF of the right leg was measured by a floor-embedded force plate (AMTI, USA) at 1,000Hz, low-pass filtered at 15Hz, and downsampled to 100Hz. Each trial contains 0.8s of data starting at the moment of jumping, corresponding to 80 time steps. We appended zeros to the end of each trial to match the input length of our transformer. We retained all the trials that were successfully collected. In total, we obtained 509 windows from 509 drop landing trials for model training and evaluation.

### C. Model Architecture

We built a transformer model to learn IMU representations during pre-training (Fig. 2(a)). The transformer’s input is a window of IMU data with 128 samples and 48 IMU axes (eight 3-axis accelerometers and eight 3-axis gyroscopes). We first normalized each IMU axis by removing the mean and scaling to unit variance. Then, the 128 samples were sliced into patches and the values of each patch were flattened. We randomly masked a portion of patches with a trainable masking patch that is updated during the training process. The masking ratio is a critical parameter in SSL, and it depends on the patch length of transformers. We performed a grid search to jointly optimize the masking ratio (6.25%, 12.5%, 12.5%, 25%, 37.5%, 50%, and 62.5%) and the patch length (1, 2, 4, and 8). After masking, we linearly mapped each patch to a vector of length 192 and added sinusoidal positional encoding [35]. Then, the data was fed into the transformer model, generating an output with dimensions identical to the input. Each output patch was linearly mapped to the original patch size (patch length *×* 48 IMU axes) to reconstruct the unmasked input data. The transformer was trained to minimize the mean squared error between the reconstructed data and the unmasked input data over the masked patches. The transformer has six self-attention blocks, with a per-block configuration of 8 attention heads, 512 feedforward units, 10% dropout, 10^*−*5^ layer normalization, and rectified linear unit (ReLU) activation function. The total number of trainable parameters is 2 million.

**Fig. 2.**
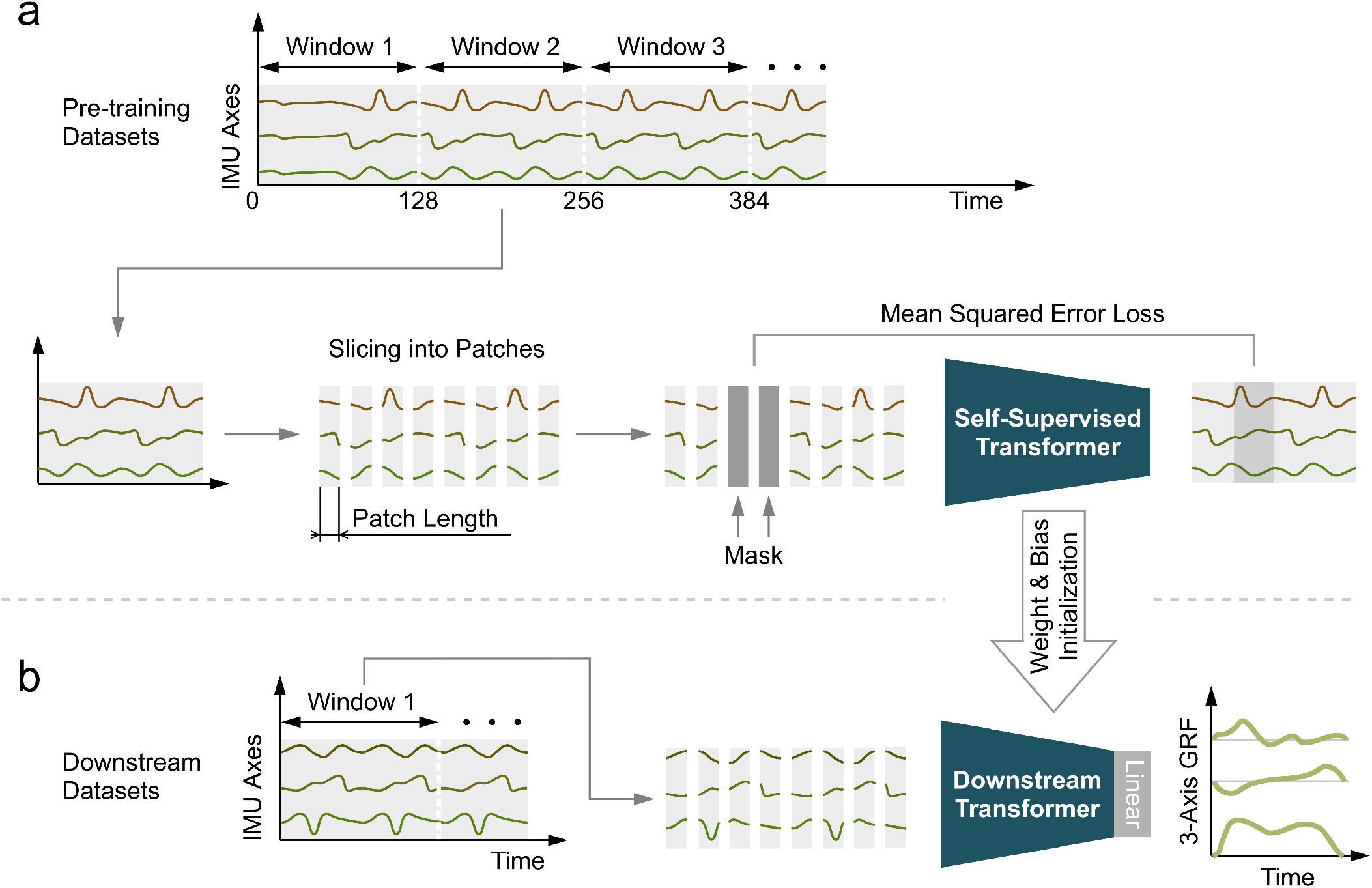
Self-supervised IMU representation learning to boost downstream prediction tasks. (a) Continuous IMU data are segmented into windows, sliced into patches, randomly masked, and finally fed into a transformer model to reconstruct the original data window. (b) The weights and biases of the self-supervised transformer were copied to initialize those of the transformer for downstream evaluation. A linear layer was appended to the transformer to map the transformer outputs to the final model outputs. During fine-tuning, we optimized the linear layer first and then fine-tuned the entire model to prevent distortion of pre-trained IMU representations [33], [34]. The fine-tuning data preprocessing steps are identical to those used in SSL, except for the exclusion of masking.

The model for downstream prediction tasks (i.e., estimating 3-axis GRF) has the same architecture as the one used in pre-training (Fig. 2(b)). Both IMU and GRF data are normalized along each axis by removing the mean and scaling to unit variance. The output of each patch was linearly mapped to GRF. The data pre-processing steps are identical to those in pre-training except that no patch is masked. For each downstream task, we investigated the performance of three pre-trained models whose parameters are inherited from SSL pre-training on the three different pre-training datasets.

The proposed model uses data from eight IMUs as input (Fig. 1). For downstream tasks that used a smaller IMU set (e.g., overground walking [25]), we substituted each omitted IMU with a randomly initialized, trainable data patch that is updated during the training process. Prior studies reported high GRF estimation accuracy using a single foot-worn IMU [36], [37]. Thus, we trained additional models using solely the IMU data from the right foot, in addition to the models trained with data from all IMUs.

### D. Baseline and Other Pre-trained Models

The following baseline and pre-trained models have an identical model architecture to the proposed SSL model. Also, for downstream evaluation, the same fine-tuning protocol was applied to all the models.

#### 1) Baseline (No Pre-Training)

Parameters of this model are randomly initialized following conventional supervised learning with PyTorch’s default settings. Specifically, we set linear layer weights following the Kaiming uniform method [38], attention layer weights following the Xavier uniform method [39], and model biases to zeros.

#### 2) Motion Transfer Pre-training

This baseline model is pre-trained on a running dataset with real IMU data as input and 3-axis GRF data as output. The distribution shift between this pre-training and downstream dataset is predominantly from the motion. The data were collected from 15 participants running on an instrumented treadmill with various speeds, footwear, strike patterns, and step rates [14]. The GRF is measured by an instrumented treadmill (Bertec, USA) at 1,000Hz and low-pass filtered at 15Hz. Five IMUs (MTi-300, Xsens Technologies B.V., the Netherlands) were placed on the trunk, pelvis, and left thigh, shank, and foot to collect data at 200Hz. We directly used the left thigh, shank, and foot IMU data to substitute the absent right thigh, shank, and foot IMU. We downsampled IMU and GRF data to 100Hz and segmented each trial into windows of 128 time steps (1.28*s* each) with 50% overlap (0.64*s*) to match those of downstream datasets. Both IMU and GRF data are normalized along each axis by removing the mean and scaling to unit variance. In total, there were 21,099 windows available for model pre-training.

#### 3) Task Transfer Pre-training

This baseline model is pre-trained on the AMASS dataset (Section II-A2) with synthetic IMU data as input and 3-axis body center acceleration in the global frame as output. The body center acceleration, which is measured by marker-based motion capture, is selected due to its association with GRF according to Newton’s second law. Both synthetic IMU data and body center acceleration are normalized along each axis by removing the mean and scaling to unit variance.

### E. Training Protocols

Models were implemented in Python 3.9 with PyTorch 1.13 and Numpy 1.23. One Nvidia RTX 2080 Ti graphics card was used to train and test all the models. For the proposed SSL pre-training, Motion Transfer Pre-training, and Task Transfer Pre-training, we used the AdamW optimizer [40] to minimize the mean squared error with a learning rate of 10^*−*4^, a weight decay of 0.01, and a minibatch size of 64 for 5 *×* 10^4^ backpropagation steps without early stopping. We used a linear warmup of the learning rate for the first 20% of steps, and decayed the learning rate with a cosine decay schedule without restarts [41]. We used the entire dataset for pre-training without splitting a test set because the model evaluation was performed based on downstream datasets that are different from pre-training datasets.

For downstream evaluation, we implemented a two-step fine-tuning strategy for pre-trained models, which optimizes the linear layer first and then fine-tunes the entire model to prevent distortion of pre-trained IMU representations [33], [34]. For the first step where the linear layer was trained, we used AdamW optimizer with a minibatch size of 64, a learning rate of 10^*−*3^, and a weight decay of 0.01 for 300 backpropagation steps. For the second step where the entire model is fine-tuned, we used AdamW optimizer with a minibatch size of 64, a learning rate of 10^*−*4^, and a weight decay of 0.01 for 300 backpropagation steps. For both steps, we used the mean squared error loss function, linearly warmed up the learning rate for the first 20% of backpropagation steps, and decayed the learning rate with a cosine decay schedule without restarts [41]. We trained and tested models with data from different participants for each dataset using a five-fold cross-validation approach, where participants were randomly split into five groups. Four groups were combined as the training set for each of the five iterations, and the remaining group was used as the test set.

### F. Performance Evaluation

We compared different models using Pearson’s correlation (*ρ*) and root mean square error (RMSE) between the estimated and gold-standard GRF across all the stance phases and landing phases of each participant. Swing phases of walking and flight phases of drop landing where vGRF is lower than 2% of the body weight are excluded from the analysis. In addition to RMSE, we compared the relative RMSE (rRMSE) of peak vGRF because its magnitude is larger during drop landing than walking. The rRMSE is defined as:

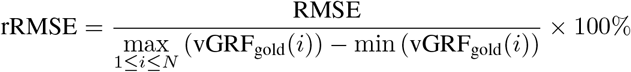

where *N* denotes the number of windows of a participant and vGRF_gold_ denotes gold-standard vGRF measurement. A oneway analysis of variance (ANOVA) was used to determine if estimation accuracy would be significantly affected by pre-training settings, i.e., baseline, using real IMU, using synthetic IMU, and using real and synthetic IMU. If significant differences were detected, paired t-tests with Bonferroni corrections were used to compare pairs of models. The level of significance was set to 0.05.

Beyond overall GRF measures, we also quantified the error of our pre-trained models across a spectrum and compared it to that of baseline models. We performed spectrum analysis using Fast Fourier Transform (FFT) on the GRF measured by the force plate as well as on the GRF estimated by models for each window. Then, we computed the differences in FFT results and averaged the differences across all the windows.

In addition, we investigated scaling law curves for model performance by varying the downstream dataset sizes. For each fold of cross-validation, we randomly sampled a subset of the training set for model training, while used the entire test set for evaluation. We linearly reduced the sizes (100%, 90%, 80%,…, 10%) for GRF estimation during overground walking and drop landing, whereas exponentially reduced the sizes (10^0^ *×* 100%, 10^*−*0.2^ *×* 100%, 10^*−*0.4^ *×* 100%, …, 10^*−*2^ *×* 100%) for GRF estimation during treadmill walking due to its larger base size.

## III. Results

SSL pre-trained models achieved higher accuracy in estimating vGRF during overground walking and treadmill walking across various downstream dataset sizes compared to baseline models that have an identical model architecture (Fig. 3). The models pre-trained on synthetic IMU data and on a combination of the real and synthetic IMU data were both significantly more accurate than the baseline models when using the entire overground or treadmill walking dataset for fine-tuning. No differences were observed between pre-trained and baseline models in estimating mlGRF, apGRF, or vGRF during drop landing. Using 10% of overground walking data to fine-tune the SSL pre-trained models yielded comparable accuracy to the baseline model trained on 100% of overground walking data, indicating a 10x data efficiency improvement. Similarly, using 1% of treadmill walking data to fine-tune the pre-trained models yielded comparable accuracy to the baseline models that are trained on 100% of treadmill walking data, indicating a 100x data efficiency improvement. There was no statistical difference between the three SSL pre-trained models for all three prediction tasks except that the model pre-trained on the synthetic IMU dataset was significantly more accurate than the model pre-trained on the real IMU dataset for treadmill walking. Apart from the baseline model, the proposed SSL achieved slightly better results than Motion Transfer, which pre-trained the model using running GRF (Table II-F). The proposed SSL achieved comparable results to Task Transfer, which pre-trained the model using body center acceleration measured by marker-based motion capture.

**TABLE II.**
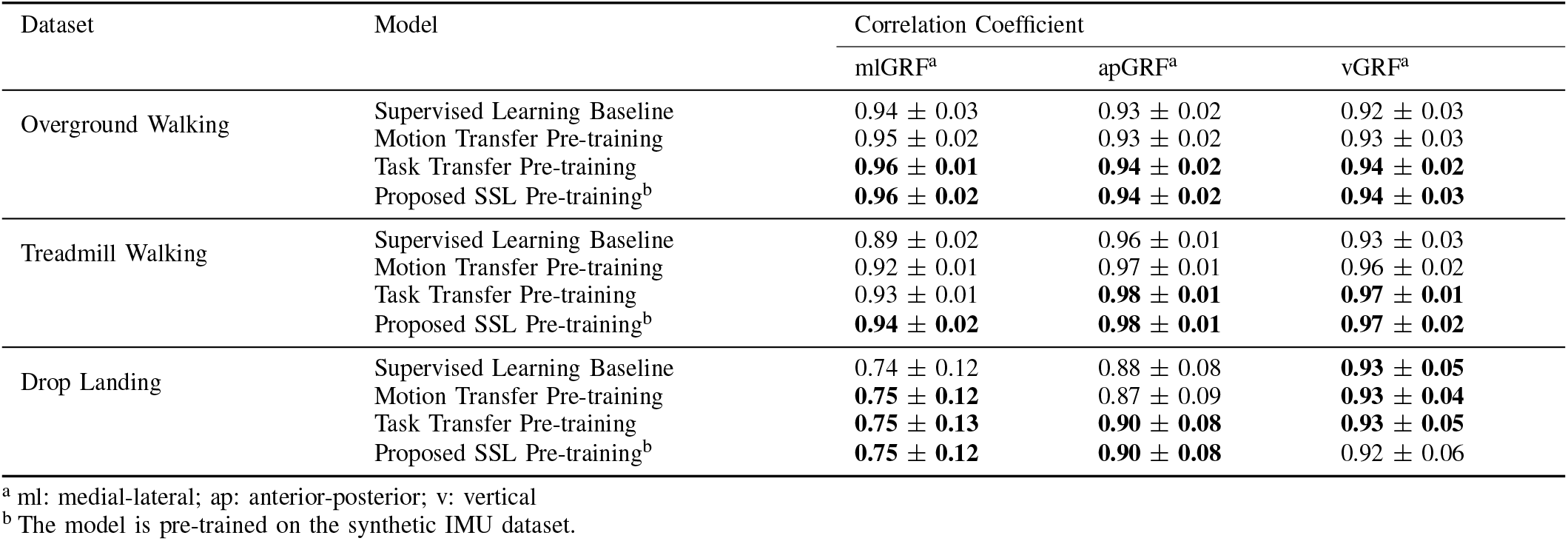
Correlation Coefficient between the Gold-Standard and Estimated GRF for the Baseline and a Variety of Pre-trained Models.

| Dataset | Model | Correlation Coefficient |  |  |
| --- | --- | --- | --- | --- |
|  |  | mlGRF<sup>a</sup> | apGRF<sup>a</sup> | vGRF<sup>a</sup> |
| Overground Walking | Supervised Learning Baseline | 0.94 ± 0.03 | 0.93 ± 0.02 | 0.92 ± 0.03 |
|  | Motion Transfer Pre-training | 0.95 ± 0.02 | 0.93 ± 0.02 | 0.93 ± 0.03 |
|  | Task Transfer Pre-training | **0.96 ± 0.01** | **0.94 ± 0.02** | **0.94 ± 0.02** |
|  | Proposed SSL Pre-training<sup>b</sup> | **0.96 ± 0.02** | **0.94 ± 0.02** | **0.94 ± 0.03** |
| Treadmill Walking | Supervised Learning Baseline | 0.89 ± 0.02 | 0.96 ± 0.01 | 0.93 ± 0.03 |
|  | Motion Transfer Pre-training | 0.92 ± 0.01 | 0.97 ± 0.01 | 0.96 ± 0.02 |
|  | Task Transfer Pre-training | 0.93 ± 0.01 | **0.98 ± 0.01** | **0.97 ± 0.01** |
|  | Proposed SSL Pre-training<sup>b</sup> | **0.94 ± 0.02** | **0.98 ± 0.01** | **0.97 ± 0.02** |
| Drop Landing | Supervised Learning Baseline | 0.74 ± 0.12 | 0.88 ± 0.08 | **0.93 ± 0.05** |
|  | Motion Transfer Pre-training | **0.75 ± 0.12** | 0.87 ± 0.09 | **0.93 ± 0.04** |
|  | Task Transfer Pre-training | **0.75 ± 0.13** | **0.90 ± 0.08** | **0.93 ± 0.05** |
|  | Proposed SSL Pre-training<sup>b</sup> | **0.75 ± 0.12** | **0.90 ± 0.08** | 0.92 ± 0.06 |

<sup>a</sup> ml: medial-lateral; ap: anterior-posterior; v: vertical
<sup>b</sup> The model is pre-trained on the synthetic IMU dataset.

**Fig. 3.**
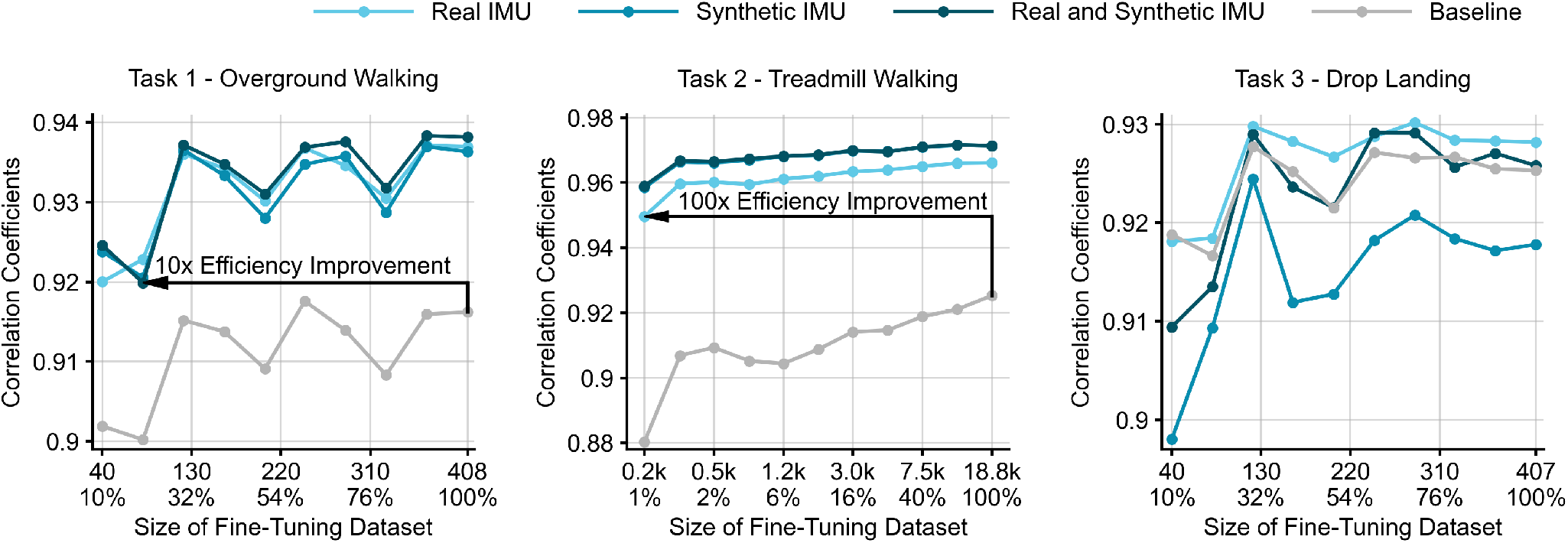
Correlation coefficient between gold-standard and estimated vGRF for SSL pre-trained and baseline models when using a range of reduced downstream datasets for training. The sizes of overground walking and drop landing datasets for model fine-tuning were reduced linearly whereas the size of treadmill walking dataset was reduced exponentially due to a significantly larger dataset size. Black arrows in task 1 - overground walking and task 2 - treadmill walking indicate improvements in data efficiency.

When evaluating performance on vGRF peaks, the SSL pre-trained models also outperformed the baseline model for overground and treadmill walking (Fig. 4). SSL pre-training increased the correlation of peak vGRF estimation from 0.59 to 0.64 for overground walking and from 0.69 to 0.90 for treadmill walking. SSL pre-training reduced the correlation of peak vGRF estimation from 0.70 to 0.68 for drop landing. The RMSEs and rRMSE of pre-trained models are 0.11±0.03*N/kg* and 8.2±2.4% for overground walking, 0.07±0.02*N/kg* and 4.5±1.0% for treadmill walking, and 0.20*±*0.06*N/kg* and 7.9*±*2.8% for drop landing, respectively.

**Fig. 4.**
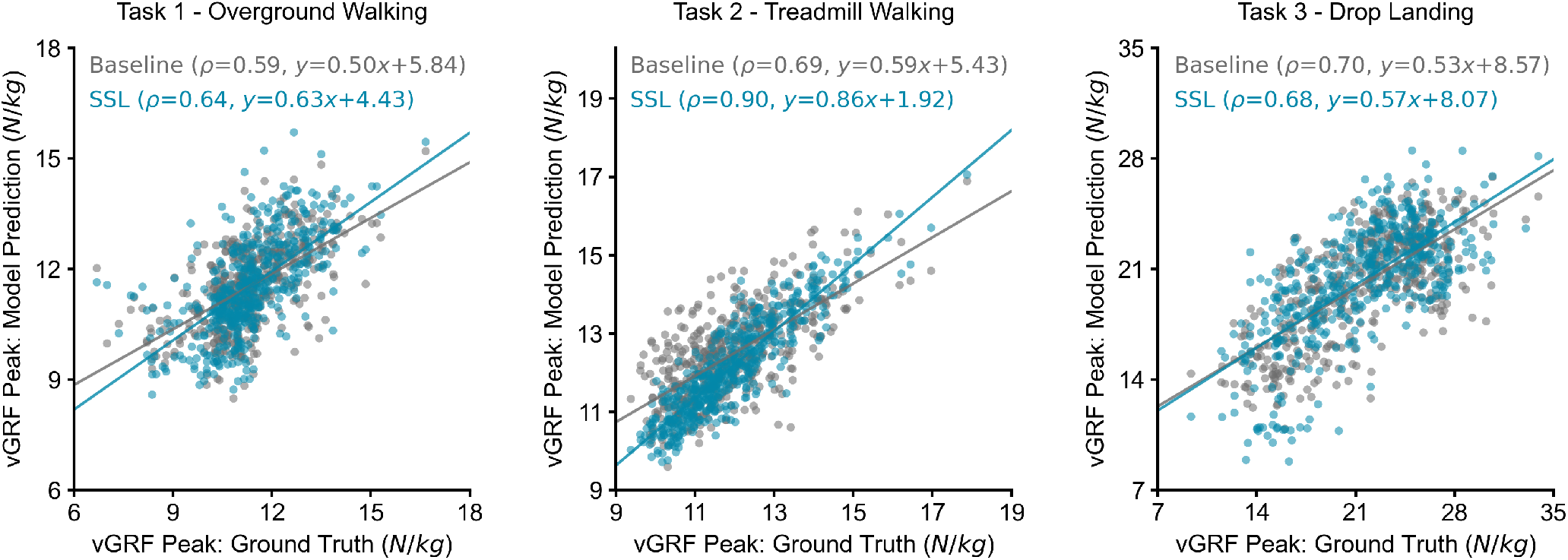
Scatter plot and regression line showing gold-standard and estimated vGRF peaks for SSL pre-trained and baseline models. All the windows are shown for task 1 (overground walking) and task 3 (drop landing), whereas 500 windows were randomly selected from 23,481 windows for task 2 (treadmill walking).

When using the right foot IMU data as model input, SSL pre-trained models consistently achieved higher accuracy than the baseline model across mlGRF, apGRF, and vGRF during overground walking, treadmill walking, and drop landing (Table III). Also, using a single foot-worn IMU resulted in lower correlation coefficients compared to using all the IMUs. Additionally, consistently smaller errors were observed in SSL pre-trained models compared to the baseline models across the spectrum for overground and treadmill walking (Fig. 5). For drop landing, the SSL pre-trained model exhibited comparable error to the baseline model across the spectrum. A masking ratio of 6.25–12.5% with shorter patch lengths achieved the best performance across all three prediction tasks (Fig. 6). Thus, we implemented our self-supervised transformer models with a patch length of 1 (equivalent to 10 ms at 100Hz sampling frequency) and 12.5% masking, and all the above results are reported based on these models.

**TABLE III.** Correlation Coefficient between the Gold-Standard and Estimated GRF with a Single Foot IMU Data as Model Input.

| Dataset | Model | Correlation Coefficient |  |  |
| --- | --- | --- | --- | --- |
|  |  | mlGRF<sup>a</sup> | apGRF<sup>a</sup> | vGRF<sup>a</sup> |
| Overground Walking | Baseline | 0.93±0.03 | 0.91±0.02 | 0.90±0.04 |
|  | Proposed SSL<sup>b</sup> | 0.94±0.03 | 0.93±0.03 | 0.93±0.03 |
| Treadmill Walking | Baseline | 0.84±0.04 | 0.93±0.02 | 0.87±0.03 |
|  | Proposed SSL<sup>b</sup> | 0.90±0.03 | 0.97±0.01 | 0.94±0.02 |
| Drop Landing | Baseline | 0.70±0.12 | 0.86±0.07 | 0.88±0.06 |
|  | Proposed SSL<sup>b</sup> | 0.72±0.12 | 0.87±0.06 | 0.89±0.06 |

<sup>a</sup> ml: medial-lateral; ap: anterior-posterior; v: vertical
<sup>b</sup> The model is pre-trained on the synthetic IMU dataset.

**Fig. 5.**
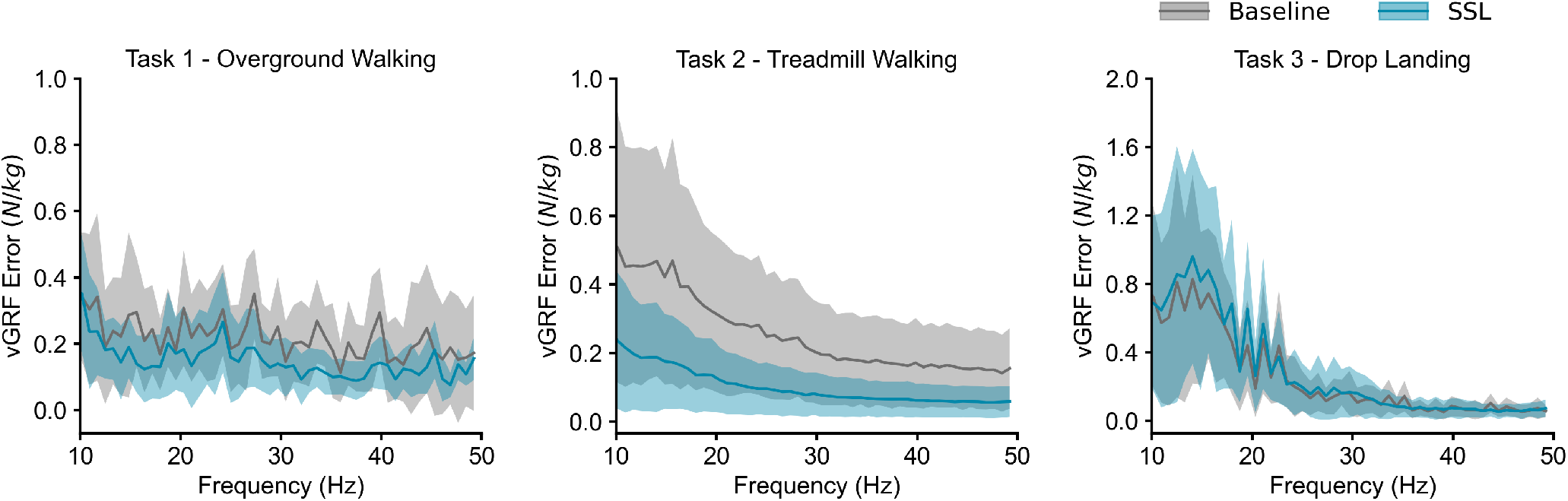
The error between gold-standard measurements and model estimations across the spectrum. Pre-trained models (blue) exhibited smaller errors compared to baseline models (gray) for overground and treadmill walking.

**Fig. 6.**
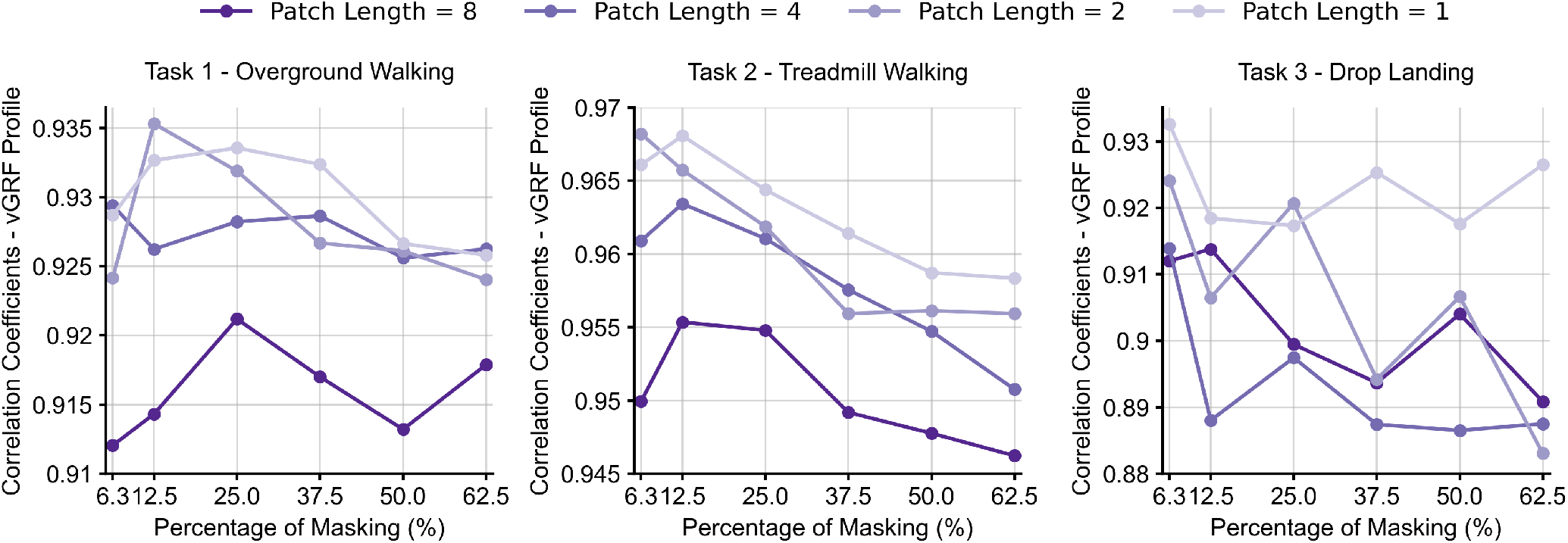
Correlation coefficient between the gold-standard and estimated vGRF for a variety of models trained with patch lengths of 1–8 and with 6.25–62.5% masking during pre-traing. Patch lengths of 1–2 and 6.25–12.5% masking have the highest accuracy.

## IV. Discussion

This work shows that using SSL to pre-train deep learning models that predict walking GRF from IMUs can improve data efficiency. Prior deep learning models for biomechanical estimation are predominantly trained on proprietary datasets with a small number of participants. In contrast, we used open-source motion datasets with hundreds of participants to pre-train our models. Three key outcomes of this work are:

- Compared to conventional supervised learning, using SSL and large “unlabeled” IMU datasets to pre-train deep learning models can improve data efficiency by 10–100x for IMU-based walking vGRF estimation. It can also improve correlation coefficient by 0.02-0.04 when using the same amount of labeled data for training.
- Low masking ratios (6.25-12.5%) are optimal for IMU-based SSL, indicating that IMU data have high information redundancy (Fig. 6).
- This code and pre-trained model have been made publicly available, potentially facilitating researchers to improve the results of their customized downstream prediction tasks (Appendix A).

In the absence of large-scale real IMU datasets, we utilized a large-scale motion dataset (AMASS) to generate synthetic IMU data for SSL. Its combined window duration is equivalent to 56 hours of data, substantially exceeding prior kinetic estimation studies that used 1.2–6 hours of real IMU data [13]– [15] and 0.2–2 hours of synthetic IMU data [42], [43]. Models pre-trained on a relatively smaller real IMU dataset and a larger synthetic IMU dataset achieved similar performance. This suggests that the size of the synthetic dataset compensated for the distribution shift between real and synthetic data that arise from soft tissue artifacts, sensor noise, and human model simplification [28]. The other two pre-training methods, Motion Transfer and Task Transfer, achieved slightly lower or comparable results to the proposed SSL pre-training. Their non-inferior performance might be attributed to the usage of gold-standard measurements from force plates and marker-based motion capture systems. The proposed SSL only relies on “unlabeled” IMU data. Thus, the results can potentially be improved and generalized to broader use cases by using “unlabeled” IMU data collected in real-life or sports environments. Ideally, the dataset should involve a wide range of motions that are related to clinical assessments and of interest to biomechanists, such as jumping, landing, and cutting.

SSL trains models to learn IMU representations specific to pre-training data. As a result, it improved GRF estimation during walking, which is one of the most common motions in both two pre-training datasets (MoVi and AMASS). In contrast, neither dataset has instances of drop landing. Consequently, the model cannot learn IMU representations of drop landing during pre-training, resulting in no enhancement during subsequent downstream evaluation. Tailoring the distribution of datasets for specific estimation tasks is a commonly used strategy for improving the performance of deep learning [44]– [46]. To apply SSL, future researchers may need to increase the similarity between the distribution of pre-training and downstream datasets. This can be achieved by adjusting the weights of instances or by collecting additional pre-training data for the motions of the downstream datasets.

The RMSE of peak vGRF estimation of the pre-trained models were 0.56–0.88*N/kg* during treadmill and overground walking, which are comparable to the smallest detectable change of peak vGRF during walking in patients with stroke or knee osteoarthritis (0.56–0.61*N/kg*) [47], [48]. The RMSE of peak vGRF estimation of the pre-trained models was 3.53*N/kg* during drop landing, which is higher than the smallest detectable change of peak vGRF during drop landing (0.55–0.63*N/kg*) [49]. Also, the performance of our pre-trained models is comparable to or slightly higher than those reported in prior studies that used the same downstream datasets for vGRF estimation. Specifically, a prior deep learning model achieved a correlation coefficient of 0.96 ± 0.01 for overground walking and 0.96±0.01 for treadmill walking [50], while our models achieved 0.95±0.02 and 0.97±0.01, respectively. Another prior deep learning model achieved a correlation coefficient of 0.92*±*0.11 [27] for vGRF estimation during drop landing, while our model achieved 0.93±0.04. Unlike prior studies, we excluded swing phase of walking and flight phases of drop landing for performance evaluation. The swing and flight phases have substantially smaller RMSE than stance and landing phases across three GRF axes, possibly because the prediction of zero GRF is inherently easier than that of positive and modulating GRF (Supplementary Table I). Thus, excluding swing and flight phases reduced correlations and increased RMSE of our models. Another reason for excluding swing phases is that prior studies used one IMU placed on foot for segmenting walking and running gait phases with less than 25 ms errors [51]–[53] Integration of such gait phase segmentation models into the GRF estimation model may obviate the need for GRF estimation during swing phase. Additionally, prior studies tailored their model architectures and hyperparameters for specific prediction tasks, and thus they might not be reused for other prediction tasks. In contrast, we demonstrated that our unified model architecture can be generalized to multiple downstream prediction tasks.

The optimal mask ratio is related to the information density of the input data. Previous language models used low masking ratios for natural language that is information-dense (e.g., 15% [22]), whereas vision models used high mask ratios for images that have high information redundancy (e.g., 75% [18], [19]). We observed that 6.26–12.5% masking ratios have the best performance (Fig. 6), indicating that IMU data are information-dense. Nevertheless, the optimal masking may increase when the IMU sampling frequencies are substantially higher than 100Hz due to increased information redundancy. Our proposed model used a window length of 128 samples (1.28*s*), which is longer than the typical duration of a walking gait cycle or a drop landing trial. However, we acknowledge that such a window length may limit the model performance on other tasks such as GRF estimation during balancing, which needs a longer window length to extend the temporal scope of the model.

Our code, pre-trained models, and tutorials are publicly available for researchers who are interested in using them for their downstream prediction tasks. Our models were trained with eight IMUs; however, they are robust to the distribution shift caused by a smaller IMU set. When using four IMUs for overground walking GRF estimation (Table II-F) or using one foot-worn IMU for both walking and drop landing GRF estimation (Table III), SSL pre-training improved accuracy across mlGRF, apGRF, and vGRF compared to no pre-training. Thus, the proposed SSL model can be used for sparse IMU configurations that are more practical for real-world measurements [31], [54], [55]. We recommend placing IMUs with their z-axes aligned with the segment surface normal, x-axes pointing left during standing, and y-axes perpendicular to the x- and z-axes following the right-hand rule (Fig. 1). For a different IMU-to-segment orientation placement configuration, the recommended configuration can typically be achieved through a post-hoc rotation of the IMU data.

## V. Conclusion

SSL with large IMU datasets for model pre-training can improve data efficiency and accuracy compared to conventional supervised learning when the distribution of the downstream dataset is within that of pre-training datasets. Future studies may collect “unlabeled” IMU data under real-life or sports environments to expand the scale and distribution of pre-training datasets, thus potentially unlocking newer and broader use cases of IMU-driven kinetic assessment where only limited gold-standard GRF measurements are available.

## Appendix A

### CODE AND DATA AVAILABILITY

The trained models are available on https://github.com/StanfordMIMI/SSL_IMU. The source code is also provided in the repository for researchers who are interested in applying SSL for pre-training specialized deep learning model architectures.

## Acknowledgment

This study was funded by the Joe and Clara Tsai Foundation through the Wu Tsai Human Performance Alliance, the U.S. National Institutes of Health (NIH) under grant R01 AR077604, R01 EB002524, R01 AR079431, and P41 EB027060, and the National Natural Science Foundation of China under grant 52250610217.

## REFERENCES

[1] J. Houck, J. Kneiss, S. V. Bukata, and J. E. Puzas, “Analysis of vertical ground reaction force variables during a sit to stand task in participants recovering from a hip fracture,” Clinical Biomechanics, vol. 26, no. 5, pp. 470–476, 2011.

[2] D. Hurwitz, A. Ryals, J. Case, J. Block, and T. Andriacchi, “The knee adduction moment during gait in subjects with knee osteoarthritis is more closely correlated with static alignment than radiographic disease severity, toe out angle and pain,” Journal of Orthopaedic Research, vol. 20, no. 1, pp. 101–107, 2002.

[3] L. Sharma, D. E. Hurwitz, E. J.-M. Thonar, J. A. Sum, M. E. Lenz, D. D. Dunlop, T. J. Schnitzer, G. Kirwan-Mellis, and T. P. Andriacchi, “Knee adduction moment, serum hyaluronan level, and disease severity in medial tibiofemoral osteoarthritis,” Arthritis & Rheumatism, vol. 41, no. 7, pp. 1233–1240, 1998.

[4] T. Miyazaki, M. Wada, H. Kawahara, M. Sato, H. Baba, and S. Shimada, “Dynamic load at baseline can predict radiographic disease progression in medial compartment knee osteoarthritis,” Annals of the Rheumatic Diseases, vol. 61, no. 7, pp. 617–622, 2002.

[5] I. Aerts, E. Cumps, E. Verhagen, J. Verschueren, and R. Meeusen, “A systematic review of different jump-landing variables in relation to injuries,” J Sports Med Phys Fitness, vol. 53, no. 5, pp. 509–519, 2013.

[6] B. Caulfield and M. Garrett, “Changes in ground reaction force during jump landing in subjects with functional instability of the ankle joint,” Clinical Biomechanics, vol. 19, no. 6, pp. 617–621, 2004.

[7] F. Porciuncula, A. V. Roto, D. Kumar, I. Davis, S. Roy, C. J. Walsh, and L. N. Awad, “Wearable movement sensors for rehabilitation: a focused review of technological and clinical advances,” PM&R, vol. 10, no. 9, pp. S220–S232, 2018.

[8] E. Dorschky, M. Nitschke, A.-K. Seifer, A. J. van den Bogert, and B. M. Eskofier, “Estimation of gait kinematics and kinetics from inertial sensor data using optimal control of musculoskeletal models,” Journal of biomechanics, vol. 95, p. 109278, 2019.

[9] T. Li, L. Wang, J. Yi, Q. Li, and T. Liu, “Reconstructing walking dynamics from two shank-mounted inertial measurement units,” IEEE/ASME Transactions on Mechatronics, vol. 26, no. 6, pp. 3040–3050, 2021.

[10] J. Kloeckner, R. Visscher, W. R. Taylor, E. Viehweger, and E. De Pieri, “Prediction of ground reaction forces and moments during walking in children with cerebral palsy,” Frontiers in Human Neuroscience, vol. 17, p. 1127613, 2023.

[11] F. De Groote and A. Falisse, “Perspective on musculoskeletal modelling and predictive simulations of human movement to assess the neuromechanics of gait,” Proceedings of the Royal Society B, vol. 288, no. 1946, p. 20202432, 2021.

[12] M. Ackermann and A. J. Van den Bogert, “Optimality principles for model-based prediction of human gait,” Journal of biomechanics, vol. 43, no. 6, pp. 1055–1060, 2010.

[13] F. J. Wouda, M. Giuberti, G. Bellusci, E. Maartens, J. Reenalda, B.-J. F. Van Beijnum, and P. H. Veltink, “Estimation of vertical ground reaction forces and sagittal knee kinematics during running using three inertial sensors,” Frontiers in Physiology, vol. 9, p. 218, 2018.

[14] T. Tan, Z. A. Strout, and P. B. Shull, “Accurate impact loading rate estimation during running via a subject-independent convolutional neural network model and optimal imu placement,” IEEE Journal of Biomedical and Health Informatics, vol. 25, no. 4, pp. 1215–1222, 2020.

[15] R. S. Alcantara, W. B. Edwards, G. Y. Millet, and A. M. Grabowski, “Predicting continuous ground reaction forces from accelerometers during uphill and downhill running: a recurrent neural network solution,” PeerJ, vol. 10, p. e12752, 2022.

[16] T. Tan, A. A. Gatti, B. Fan, K. G. Shea, S. L. Sherman, S. D. Uhlrich, J. L. Hicks, S. L. Delp, P. B. Shull, and A. S. Chaudhari, “A scoping review of portable sensing for out-of-lab anterior cruciate ligament injury prevention and rehabilitation,” NPJ Digital Medicine, vol. 6, no. 1, p. 46, 2023.

[17] T. Brown, B. Mann, N. Ryder, M. Subbiah, J. D. Kaplan, P. Dhariwal Neelakantan, P. Shyam, G. Sastry, A. Askell et al., “Language models are few-shot learners,” Advances in Neural Information Processing Systems, vol. 33, pp. 1877–1901, 2020.

[18] K. He, X. Chen, S. Xie, Y. Li, P. Dollar, and R. Girshick, “Masked autoencoders are scalable vision learners,” in Proceedings of the IEEE/CVF conference on computer vision and pattern recognition, 2022, pp. 16 000–16 009.

[19] Y. Li, H. Fan, R. Hu, C. Feichtenhofer, and K. He, “Scaling language-image pre-training via masking,” in Proceedings of the IEEE/CVF Conference on Computer Vision and Pattern Recognition, 2023, pp. 23 390–23 400.

[20] D. Hendrycks, M. Mazeika, S. Kadavath, and D. Song, “Using selfsupervised learning can improve model robustness and uncertainty,” Advances in Neural Information Processing Systems, vol. 32, 2019.

[21] Y. Sun, X. Wang, Z. Liu, J. Miller, A. Efros, and M. Hardt, “Test-time training with self-supervision for generalization under distribution shifts,” in International Conference on Machine Learning, 2020, pp. 9229–9248.

[22] J. Devlin, M.-W. Chang, K. Lee, and K. Toutanova, “BERT: Pretraining of deep bidirectional transformers for language understanding,” in Proceedings of the 2019 Conference of the North American Chapter of the Association for Computational Linguistics: Human Language Technologies, Jun. 2019, pp. 4171–4186.

[23] S. Ghorbani, K. Mahdaviani, A. Thaler, K. Kording, D. J. Cook, G. Blohm, and N. F. Troje, “Movi: A large multi-purpose human motion and video dataset,” Plos one, vol. 16, no. 6, p. e0253157, 2021.

[24] N. Mahmood, N. Ghorbani, N. F. Troje, G. Pons-Moll, and M. J. Black, “Amass: Archive of motion capture as surface shapes,” in Proceedings of the IEEE/CVF International Conference on Computer Vision, 2019, pp. 5442–5451.

[25] J. Camargo, A. Ramanathan, W. Flanagan, and A. Young, “A comprehensive, open-source dataset of lower limb biomechanics in multiple conditions of stairs, ramps, and level-ground ambulation and transitions,” Journal of Biomechanics, vol. 119, p. 110320, 2021.

[26] T. Tan, D. Wang, P. B. Shull, and E. Halilaj, “Imu and smartphone camera fusion for knee adduction and knee flexion moment estimation during walking,” IEEE Transactions on Industrial Informatics, vol. 19, no. 2, pp. 1445–1455, 2022.

[27] T. Sun, D. Li, B. Fan, T. Tan, and P. B. Shull, “Real-time ground reaction force and knee extension moment estimation during drop landings via modular lstm modeling and wearable imus,” IEEE Journal of Biomedical and Health Informatics, 2023.

[28] T. Zimmermann, B. Taetz, and G. Bleser, “Imu-to-segment assignment and orientation alignment for the lower body using deep learning,” Sensors, vol. 18, no. 1, p. 302, 2018.

[29] T. Tan, D. P. Chiasson, H. Hu, and P. B. Shull, “Influence of imu position and orientation placement errors on ground reaction force estimation,” Journal of Biomechanics, vol. 97, p. 109416, 2019.

[30] E. Rapp, S. Shin, W. Thomsen, R. Ferber, and E. Halilaj, “Estimation of kinematics from inertial measurement units using a combined deep learning and optimization framework,” Journal of Biomechanics, vol. 116, p. 110229, 2021.

[31] Y. Jiang, Y. Ye, D. Gopinath, J. Won, A. W. Winkler, and C. K. Liu, “Transformer inertial poser: Real-time human motion reconstruction from sparse imus with simultaneous terrain generation,” in SIGGRAPH Asia 2022 Conference Papers, 2022, pp. 1–9.

[32] X. Yi, Y. Zhou, and F. Xu, “Transpose: Real-time 3d human translation and pose estimation with six inertial sensors,” ACM Transactions on Graphics, vol. 40, no. 4, Aug. 2021.

[33] A. Kumar, A. Raghunathan, R. Jones, T. Ma, and P. Liang, “Fine-tuning can distort pretrained features and underperform out-of-distribution,” in International Conference on Learning Representations, 2022.

[34] J. Dominic, N. Bhaskhar, A. D. Desai, A. Schmidt, E. Rubin, B. Gunel, G. E. Gold, B. A. Hargreaves, L. Lenchik, R. Boutin et al., “Improving data-efficiency and robustness of medical imaging segmentation using inpainting-based self-supervised learning,” Bioengineering, vol. 10, no. 2, p. 207, 2023.

[35] A. Vaswani, N. Shazeer, N. Parmar, J. Uszkoreit, L. Jones, A. N. Gomez, Ł. Kaiser, and I. Polosukhin, “Attention is all you need,” Advances in Neural Information Processing Systems, vol. 30, 2017.

[36] X. Jiang, C. Napier, B. Hannigan, J. J. Eng, and C. Menon, “Estimating vertical ground reaction force during walking using a single inertial sensor,” Sensors, vol. 20, no. 15, p. 4345, 2020.

[37] M. I. M. Refai, B.-J. F. Van Beijnum, J. H. Buurke, and P. H. Veltink, “Portable gait lab: estimating 3d grf using a pelvis imu in a foot imu defined frame,” IEEE transactions on neural systems and rehabilitation engineering, vol. 28, no. 6, pp. 1308–1316, 2020.

[38] K. He, X. Zhang, S. Ren, and J. Sun, “Delving deep into rectifiers: Surpassing human-level performance on imagenet classification,” in Proceedings of the IEEE International Conference on Computer Vision, 2015, pp. 1026–1034.

[39] X. Glorot and Y. Bengio, “Understanding the difficulty of training deep feedforward neural networks,” in Proceedings of the Thirteenth International Conference on Artificial Intelligence and Statistics. JMLR Workshop and Conference Proceedings, 2010, pp. 249–256.

[40] I. Loshchilov and F. Hutter, “Decoupled weight decay regularization,” in International Conference on Learning Representations, 2018.

[41] I. Loshchilov and F. Hutter, “Sgdr: Stochastic gradient descent with warm restarts,” in International Conference on Learning Representations, 2016.

[42] M. Mundt, A. Koeppe, S. David, T. Witter, F. Bamer, W. Potthast, and B. Markert, “Estimation of gait mechanics based on simulated and measured imu data using an artificial neural network,” Frontiers in Bioengineering and Biotechnology, vol. 8, p. 41, 2020.

[43] E. Dorschky, M. Nitschke, C. F. Martindale, A. J. van den Bogert, A. D. Koelewijn, and B. M. Eskofier, “Cnn-based estimation of sagittal plane walking and running biomechanics from measured and simulated inertial sensor data,” Frontiers in Bioengineering and Biotechnology, vol. 8, p. 604, 2020.

[44] I. Ktena, O. Wiles, I. Albuquerque, S.-A. Rebuffi, R. Tanno, A. G. Roy, S. Azizi, D. Belgrave, P. Kohli, A. Karthikesalingam et al., “Generative models improve fairness of medical classifiers under distribution shifts,” arXiv preprint arXiv:2304.09218, 2023.

[45] P. Chambon, C. Bluethgen, J.-B. Delbrouck, R. Van der Sluijs, M. Połacin, J. M. Z. Chaves, T. M. Abraham, S. Purohit, C. P. Langlotz, and A. Chaudhari, “Roentgen: vision-language foundation model for chest x-ray generation,” arXiv preprint arXiv:2211.12737, 2022.

[46] Y. Meng, M. Michalski, J. Huang, Y. Zhang, T. Abdelzaher, and J. Han, “Tuning language models as training data generators for augmentation-enhanced few-shot learning,” in International Conference on Machine Learning, 2023, pp. 24 457–24 477.

[47] I. Campanini and A. Merlo, “Reliabilty, smallest real difference and concurrent validity of indices computed from grf components in gait of stroke patients,” Gait & posture, vol. 30, no. 2, pp. 127–131, 2009.

[48] D. Pacifico, R. Visscher, R. List, J. F. Item-Glatthorn, N. C. Casartelli, and N. A. Maffiuletti, “Discriminant validity and reproducibility of spatiotemporal and kinetic parameters during treadmill walking in patients with knee osteoarthritis,” Gait & posture, vol. 80, pp. 77–79, 2020.

[49] L. Howe, J. North, M. Waldron, and T. Bampouras, “Reliability of independent kinetic variables and measures of inter-limb asymmetry associated with bilateral drop-landing performance,” International Journal of Physical Education, Fitness and Sports, vol. 7, no. 3, pp. 32–47, 2018.

[50] M. S. B. Hossain, Z. Guo, and H. Choi, “Estimation of lower extremity joint moments and 3d ground reaction forces using imu sensors in multiple walking conditions: A deep learning approach,” IEEE Journal of Biomedical and Health Informatics, 2023.

[51] P. Felix, J. Figueiredo, C. P. Santos, and J. C. Moreno, “Adaptive realtime tool for human gait event detection using a wearable gyroscope,” in Human-Centric Robotics: Proceedings of CLAWAR 2017: 20th International Conference on Climbing and Walking Robots and the Support Technologies for Mobile Machines. World Scientific, 2018, pp. 653–660.

[52] M. Falbriard, F. Meyer, B. Mariani, G. P. Millet, and K. Aminian, “Accurate estimation of running temporal parameters using foot-worn inertial sensors,” Frontiers in physiology, p. 610, 2018.

[53] T. Tan, Z. A. Strout, R. T. Cheung, and P. B. Shull, “Strike index estimation using a convolutional neural network with a single, shoe-mounted inertial sensor,” Journal of Biomechanics, vol. 139, p. 111145, 2022.

[54] M. Lueken, J. Wenner, S. Leonhardt, and C. Ngo, “Using synthesized imu data to train a long-short term memory-based neural network for unobtrusive gait analysis with a sparse sensor setup,” in 2022 44th Annual International Conference of the IEEE Engineering in Medicine & Biology Society, 2022, pp. 3653–3656.

[55] T. Van Wouwe, S. Lee, A. Falisse, S. Delp, and C. K. Liu, “Diffusion inertial poser: Human motion reconstruction from arbitrary sparse imu configurations,” arXiv preprint arXiv:2308.16682, 2023.

