## Supplementary Table 1 for "Self-Supervised Learning Improves Accuracy and Data Efficiency for IMU-Based Ground Reaction Force Estimation"

Supplementary Table I RMSE between the Gold-Standard and Estimated GRF for SSL Pre-trained Model during Stance, Swing Phases of Walking and Landing, Flight Phases of Drop Landing.

| | | RMSE (*N/kg*) | | |
|---|---|---|---|---|
| Dataset | Phase | mlGRF[a] | apGRF[a] | vGRF[a] |
| Overground Walking | Stance | 0.04±0.01 | 0.03±0.01 | 0.13±0.03 |
| | Swing | 0.01±0.01 | 0.01±0.00 | 0.05±0.02 |
| Treadmill Walking | Stance | 0.02±0.00 | 0.03±0.01 | 0.12±0.02 |
| | Swing | 0.01±0.00 | 0.01±0.00 | 0.06±0.01 |
| Drop Landing | Landing | 0.04±0.01 | 0.05±0.01 | 0.19±0.05 |
| | Flight | 0.01±0.00 | 0.01±0.00 | 0.05±0.02 |

[a] ml: medial-lateral; ap: anterior-posterior; v: vertical
